# Direct intracellular visualization of Ebola virus-receptor interaction by *in situ* proximity ligation

**DOI:** 10.1101/2020.05.05.080085

**Authors:** Eva Mittler, Tanwee P Alkutkar, Rohit K Jangra, Kartik Chandran

**Author notes:** Address correspondence to Kartik Chandran. Department of Integrative Structural and Computational Biology, The Scripps Research Institute, La Jolla, California, USA.

## Abstract

Ebola virus (EBOV) entry into host cells comprises stepwise and extensive interactions of the sole viral surface glycoprotein GP with multiple host factors. During the intricate process, following virus uptake and trafficking to late endosomal/lysosomal compartments, GP is proteolytically processed to GP_CL_ by the endosomal proteases cathepsin B and L unmasking GP’s receptor-binding site. Engagement of GP_CL_ with the universal filoviral intracellular receptor Niemann-Pick C1 (NPC1) eventually culminates in fusion between viral and cellular membranes, cytoplasmic escape of the viral nucleocapsid and subsequent infection. Mechanistic delineation of the indispensable GP_CL_:NPC1 binding step has been severely hampered by the unavailability of a robust cell-based assay assessing interaction of GP_CL_ with full-length endosomal NPC1.

Here, we describe a novel *in situ* assay to monitor GP_CL_:NPC1 engagement in intact, infected cells. Visualization of the subcellular localization of binding complexes is based on the principle of DNA-assisted, antibody-mediated proximity ligation. Virus-receptor binding monitored by proximity ligation was contingent on GP’s proteolytic cleavage, and was sensitive to perturbations in the GP_CL_:NPC1 interface. Our assay also specifically decoupled detection of virus-receptor binding from steps post-receptor binding, such as membrane fusion and infection. Testing of multiple FDA-approved small molecule inhibitors revealed that drug treatments inhibited virus entry and GP_CL_:NPC1 recognition by distinctive mechanisms. Together, here we present a newly established proximity ligation assay, which will allow us to dissect cellular and viral requirements for filovirus-receptor binding, and to delineate the mechanisms of action of inhibitors on filovirus entry in a cell-based system.

**IMPORTANCE:** Ebola virus causes episodic but increasingly frequent outbreaks of severe disease in Middle Africa, as shown by a currently ongoing outbreak in the Democratic Republic of Congo. Despite considerable effort, FDA-approved anti-filoviral therapeutics or targeted interventions are not available yet. Virus host-cell invasion represents an attractive target for antivirals; however our understanding of the inhibitory mechanisms of novel therapeutics is often hampered by fragmented knowledge of the filovirus-host molecular interactions required for viral infection. To help close this critical knowledge gap, here, we report an *in situ* assay to monitor binding of the EBOV glycoprotein to its receptor NPC1 in intact, infected cells. We demonstrate that our *in situ* assay based on proximity ligation represents a powerful tool to delineate receptor-viral glycoprotein interactions. Similar assays can be utilized to examine receptor interactions of diverse viral surface proteins whose studies have been hampered until now by the lack of robust *in situ* assays.

## INTRODUCTION

Members of the family *Filoviridae*, including Ebola virus (EBOV), are emerging zoonotic pathogens that cause episodic but increasingly frequent outbreaks of a highly lethal disease in Middle Africa (1). Along with the big EBOV outbreak of 2014-16 in West Africa, the ongoing outbreak of EBOV disease in the Democratic Republic of Congo, the second largest outbreak on record with >3,400 confirmed cases and >2,200 deaths (as of April 2020) and spill-over cases to neighboring Uganda highlight the potential of filoviruses to cause health emergencies of international scope (2). There is an urgent need for effective countermeasures; however, their development is hindered by our limited understanding of filovirus-host molecular interactions required for viral entry and infection.

The single filovirus-encoded spike glycoprotein (GP) is necessary and sufficient to mediate all steps in viral entry into host cells, culminating in cytoplasmic escape of the viral nucleocapsid. Following endocytosis, virions traffic to late endosomes/lysosomes (LE/LY) (3–6), where GP gains access to multiple essential host factors. GP is proteolytically processed by endosomal cysteine cathepsins B and L (CatB and CatL, respectively), which remove the heavily glycosylated C–terminal glycan cap and mucin domain sequences in the GP1 subunit (7–9), thereby exposing a recessed receptor-binding site (RBS). This cleaved GP species (GP_CL_) binds domain C of Niemann-Pick C1 (NPC1), an ubiquitously expressed cholesterol transporter embedded in endo/lysosomal membranes that acts as an universal intracellular receptor for all filoviruses (10–14). Although GP_CL_:NPC1 recognition is a prerequisite for downstream steps in virus entry and infection, *in vitro* work suggests that NPC1 binding is not sufficient to trigger large-scale conformational changes in GP or to initiate a subsequent merger of viral and host membranes (15, 16). Indeed, NPC1’s precise role beyond GP binding, which presumably provides a physical link between virus particles and host membranes, remains elusive to date.

Because a robust cell-based assay assessing the interaction of GP_CL_ with full-length endosomal NPC1 in its native context has been unavailable, mechanistic studies of this indispensable virus-receptor interaction have been largely limited to *in vitro* assays. These assays are predominantly based on a truncated, soluble form of a single domain in NPC1, domain C, as well as on *in vitro*-cleaved GPs (11, 12). However, studies with NPC1-targeting inhibitors suggest that these assays do not fully recapitulate the authentic interaction(s) between *in situ*-cleaved GP and full-length NPC1 in its membrane context within late endosomes and/or lysosomes (16). To address this gap, we describe herein an *in situ* assay to monitor GP_CL_:NPC1 binding in individual endosomal compartments of intact, infected cells by using DNA-guided, antibody-mediated proximity ligation. We employed this assay to show that GP_CL_:NPC1 interaction is restricted to the lumina of NPC1 ^+^ LE/LY, is contingent on the proteolytic cleavage of GP and is sensitive to the mutational disruption of the GP_CL_:NPC1 interface. Testing of multiple FDA-approved small molecule inhibitors in our assay revealed that drug treatments inhibited virus entry and GP_CL_:NPC1 recognition by distinct mechanisms. Application of this assay will allow us to dissect the cellular and viral requirements for filovirus-receptor interaction, and to delineate the mechanisms of action of small molecule inhibitors on filovirus entry.

## RESULTS

### Development of an assay visualizing EBOV GP:NPC1 binding in intact cells by *in situ* proximity ligation

During viral entry, proteolytically cleaved forms of EBOV GP (GP_CL_) interact with their critical endosomal receptor NPC1. We postulated that an *in situ* proximity ligation assay (PLA) could be used to monitor this essential binding step in intact cells. To detect viral particles, we used a recombinant vesicular stomatitis virus (rVSV) containing the viral phosphoprotein P linked to a fluorescent monomeric NeonGreen (mNG-P) protein, and bearing EBOV GP (17). Viral particles were allowed to attach at 4°C to U2OS human osteosarcoma cells stably expressing NPC1 tagged with a blue fluorophore, eBFP2 (17), and the cells were then shifted to 37°C to allow synchronized viral internalization. Visualization of fixed U2OS^NPC1-eBFP2^ cells by fluorescence microscopy revealed that most internalized viral particles reached NPC1-containing late endosomes (NPC1^+^ LE) within 60 min (Fig. 1).

**Fig. 1.**
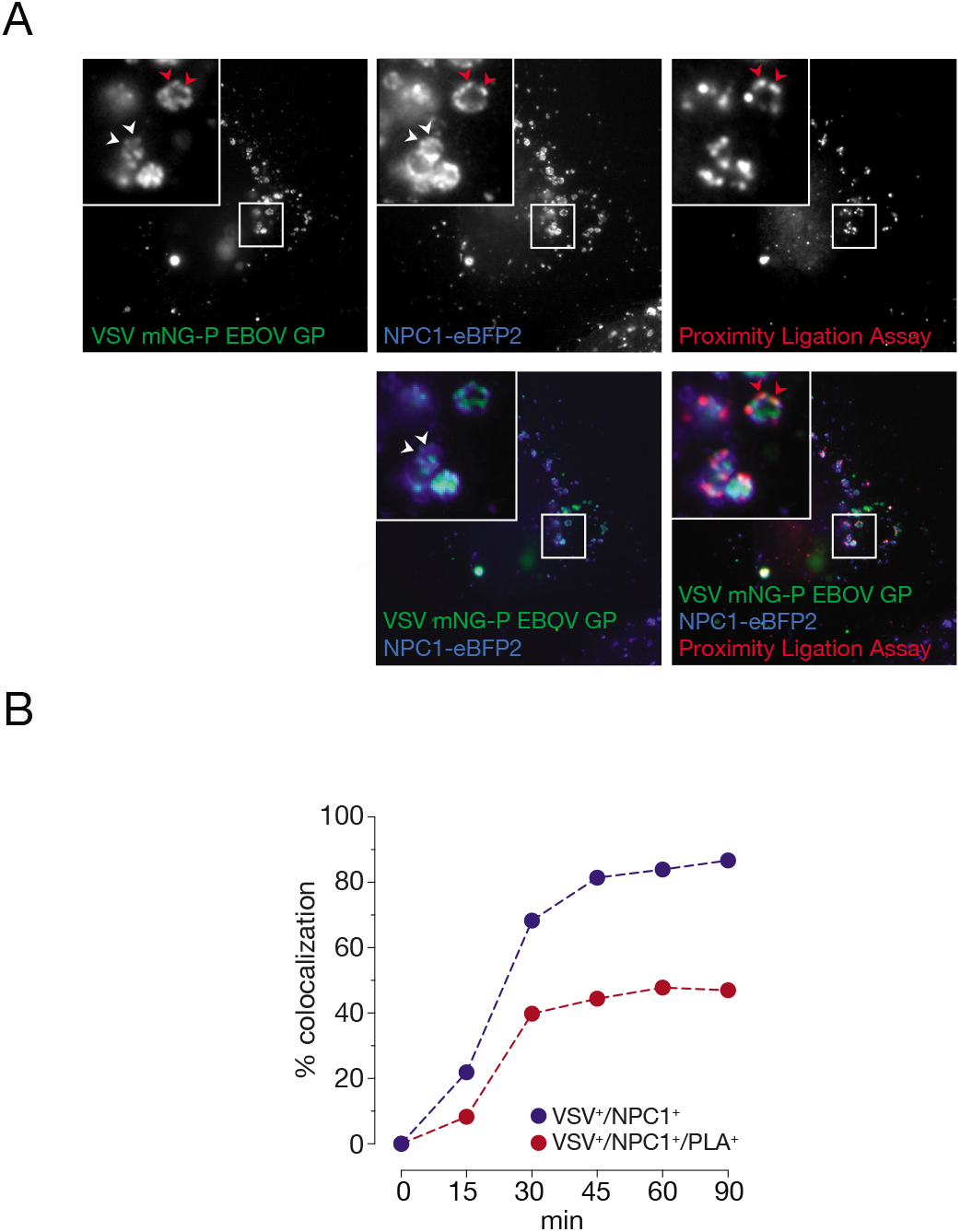
Development of an EBOV GP:NPC1 binding assay in intact cells by *in situ* proximity ligation. (**A**) VSV mNG-P particles bearing EBOV GP were internalized into U2OS cells ectopically expressing NPC1-eBFP2 for 60 min. Cells were fixed, permeabilized and subjected to proximity ligation assay (PLA) using GP_CL_- and NPC1-specific antibodies (MR72/mAb-548). During amplification, resulting PLA products were labeled with a detector oligonucleotide conjugated with a red fluorophore. White arrowheads, VSV/NPC1 colocalization; red arrowheads, VSV/NPC1/PLA colocalization. (**B**) VSVs bearing EBOV GP were endocytosed into U2OS cells as described in (A), fixed at the indicated time points and subjected to PLA. VSV positive compartments (VSV^+^) were enumerated; the percentage of compartments also positive for NPC1 (VSV^+^/NPC1^+^) or NPC1 plus PLA (VSV^+^/NPC1^+^/PLA^+^) were determined. Representative curves out of two independent experiments are shown; averages of pooled cells are displayed (*n* ≥ 20 per time point).

To detect closely apposed GP_CL_ and NPC1 molecules in infected cells, we incubated them with oligonucleotide-linked monoclonal antibodies directed against each protein: GP’s highly conserved receptor-binding site (RBS), unmasked by proteolytic cleavage to GP_CL_ in endosomes, was detected with the RBS-specific antibody MR72 (17–19), and NPC1 was detected with the domain C-specific antibody mAb-548 (17). Circular DNA molecules were generated by oligonucleotide-guided proximity ligation, amplified *in situ* and visualized with a fluorophore-conjugated detector oligonucleotide (20, 21). PLA signal required the addition of both, MR72 and mAb-548 antibodies, and was only observed in cells that were allowed to internalize viral particles (Fig. S1). Also, PLA signal colocalized with VSV mNG-P particles bearing EBOV GP in the lumina of NPC1^+^ LE (Fig. 1A). Proximity ligation and viral trafficking to NPC1^+^ LE displayed similar kinetics—both peaked within 60 min post-viral uptake, followed by a plateau phase (Figs. 1A and B). Although most internalized viral particles trafficked to and colocalized with NPC1^+^ compartments (Fig. 1B), only a subset of these VSV^+^/NPC1^+^ vesicles also displayed PLA signal (Fig. 1B). Accordingly, in the following studies aimed at determining the viral and cellular requirements for proximity-dependent ligation of GP_CL^-^_ and NPC1-specific antibodies, we routinely enumerated the number of VSV^+^/NPC1^+^ compartments per cell, and reported the percentage of these compartments that were also positive for PLA signal (VSV^+^/NPC1^+^/PLA^+^).

### Proximity ligation is sensitive to perturbations in the GP_CL_:NPCI domain C interface

Specific interactions of EBOV GP with NPC1 domain C have been mapped to amino acid residues in the GP1 subunit that are exposed upon proteolytic cleavage (11, 18, 22). Mutations of the charged surface-exposed amino acids K114/K115 and the polar amino acid T83 led to significant defects in NPC1 binding and viral entry (5, 18). To determine if these mutations impact proximity ligation, we exposed cells to VSVs bearing GP^T83M/K114E/K115E^. Virions were efficiently trafficked to NPC1^+^ LE and underwent GP1 cleavage, as indicated by detection of the GP1 RBS with the mAb MR72. However, viral infectivity was greatly diminished (Figs. S2A and S2B). Concordantly, we observed significant reduction of VSV^+^/NPC1^+^/PLA^+^ colocalization, suggesting that GP_CL_:NPC1 engagement in LE is necessary for PLA signal formation (Fig. 2A).

**Fig. 2.**
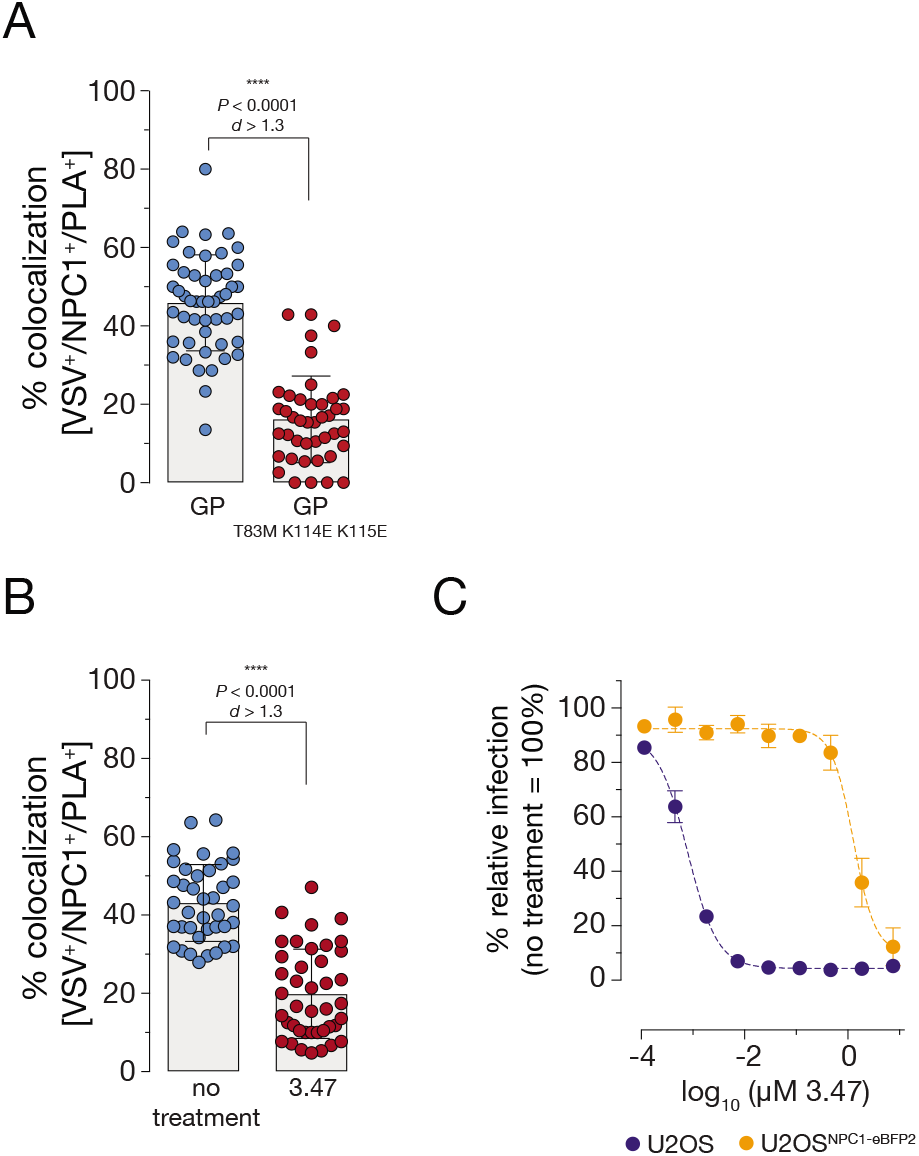
Proximity ligation is sensitive to perturbations in the GP_CL_:NPC1 domain C interface. (**A**) VSV mNG-P particles bearing EBOV GP or EBOV GP^T83M/K114E/K115E^ were internalized into U2OS^NPC1-eBFP2^ cells for 60 min followed by cell fixation, permeabilization and PLA. Cells were analyzed by fluorescence microscopy; data points represent the percentage of VSV^+^/NPC1^+^/PLA^+^ compartments per individual cell; bars depict the pooled averages and standard deviations (± SD) for all cells from two independent experiments (*n* ≥ 40). An unpaired two-tailed t-test was used to compare VSV^+^/NPC1^+^/PLA^+^ compartments of cells infected by VSV bearing either EBOV GP or EBOV GP^T83M/K114E/K115E^ (****, *P* < 0.0001). Group means calculated from the percentage of VSV^+^/NPC1^+^/PLA^+^ vesicles were compared by Cohen’s *d* effect size (*d* > 1.3). (**B**) U2OS^NPC1-eBFP2^ cells were pre-incubated with the inhibitor 3.47 (1 μM) for 60 min at 37°C, followed by VSV mNG-P EBOV GP uptake for 60 min and PLA. Data points were acquired and analyzed as described in (A). (**C**) After pre-incubation of U2OS^NPC1-eBFP2^ and wild-type U2OS cells with increasing 3.47 concentrations, cells were infected with VSV mNG-P EBOV GP for 16 h. Infection was measured by automated counting of mNG^+^ cells and normalized to infection obtained in the absence of 3.47. Averages ± SD for six technical replicates pooled from two independent experiments are displayed. Data was subjected to nonlinear regression analysis to derive 3.47 concentration at half-maximal inhibition of infection (IC_50_ ± 95% confidence intervals for nonlinear curve fit).

The small molecule filovirus entry inhibitor 3.47 was described to block EBOV GP-dependent entry and infection potentially by interfering with GP_CL_:NPC1 recognition (12). Accordingly, we directly compared the effect of 3.47 on viral entry in wild-type U2OS cells expressing basal levels of NPC1 against that in U2OS^NPC1-eBFP2^ cells used for the PLA (Fig. 2C). Consistent with its activity as an NPC1-targeting inhibitor, and as shown previously, 3.47’s antiviral activity was substantially attenuated in cells over-expressing NPC1, but we were nevertheless able to identify a concentration (1 μM) that afforded ~50% inhibition of viral entry; higher concentrations of 3.47 were cytotoxic. Pre-treatment of U2OS^NPC1-eBFP2^ cells with 3.47 at 1 μM significantly reduced PLA signal (Fig. 2B) but did not modify NPC1^+^ LE morphology, block viral trafficking and GP cleavage, or influence detection by the assay antibodies, MR72 and mAb-548 (Fig. S2C). Our data that both genetic and pharmacological disruptions of the virus-receptor interface inhibit PLA strongly suggests that this assay indeed monitors GP_CL_:NPC1 domain C engagement in endosomal compartments.

### Detection of GP_CL_:NPC1 binding by PLA requires endosomal cleavage of GP

Endosomal host cysteine cathepsins B and L (CatB and CatL, respectively) are key mediators of the entry-related GP→GP_CL_ cleavage that is required for GP_CL_:NPC1 binding and viral membrane fusion (5, 7–9). Their inactivation by pan-cysteine cathepsin inhibitors, including E-64 and E-64d, blocks filovirus entry ((7, 8); also see Fig. S3A). Concordantly, we found that pre-treatment of cells with E-64d abolished both GP cleavage (and consequently exposure of the GP1 NPC1-binding site recognized by MR72; Fig. 3B) and GP_CL_:NPC1 engagement as measured by PLA (Fig. 3A). Intriguingly, previous work has also hinted at the existence of cellular target(s) of E-64 in addition to CatB and CatL, whose inhibition imposes one or more blocks to viral entry (7, 8, 23, 24). To further investigate the cysteine protease-dependent EBOV entry mechanism, we generated *CatB/L*-knockout (KO) U2OS cell lines by CRISPR/Cas9 genome engineering (Fig. S4A). As expected, the *CatB/L-KO* cells lacked CatB and CatL activity and were substantially resistant to EBOV GP-dependent entry (Fig. S4B and C). Surprisingly, however, and in contrast to our findings in E-64d treated cells, viral particles underwent efficient exposure of the NPC1-binding site (and MR72 epitope) in GP in *CatB/L*-KO cells (Fig. 3B). Despite this apparent capacity for GP cleavage in the absence of CatB and CatL, viral particles were nevertheless unable to engage NPC1 as measured by PLA (Fig. 3A).

**Fig. 3.**
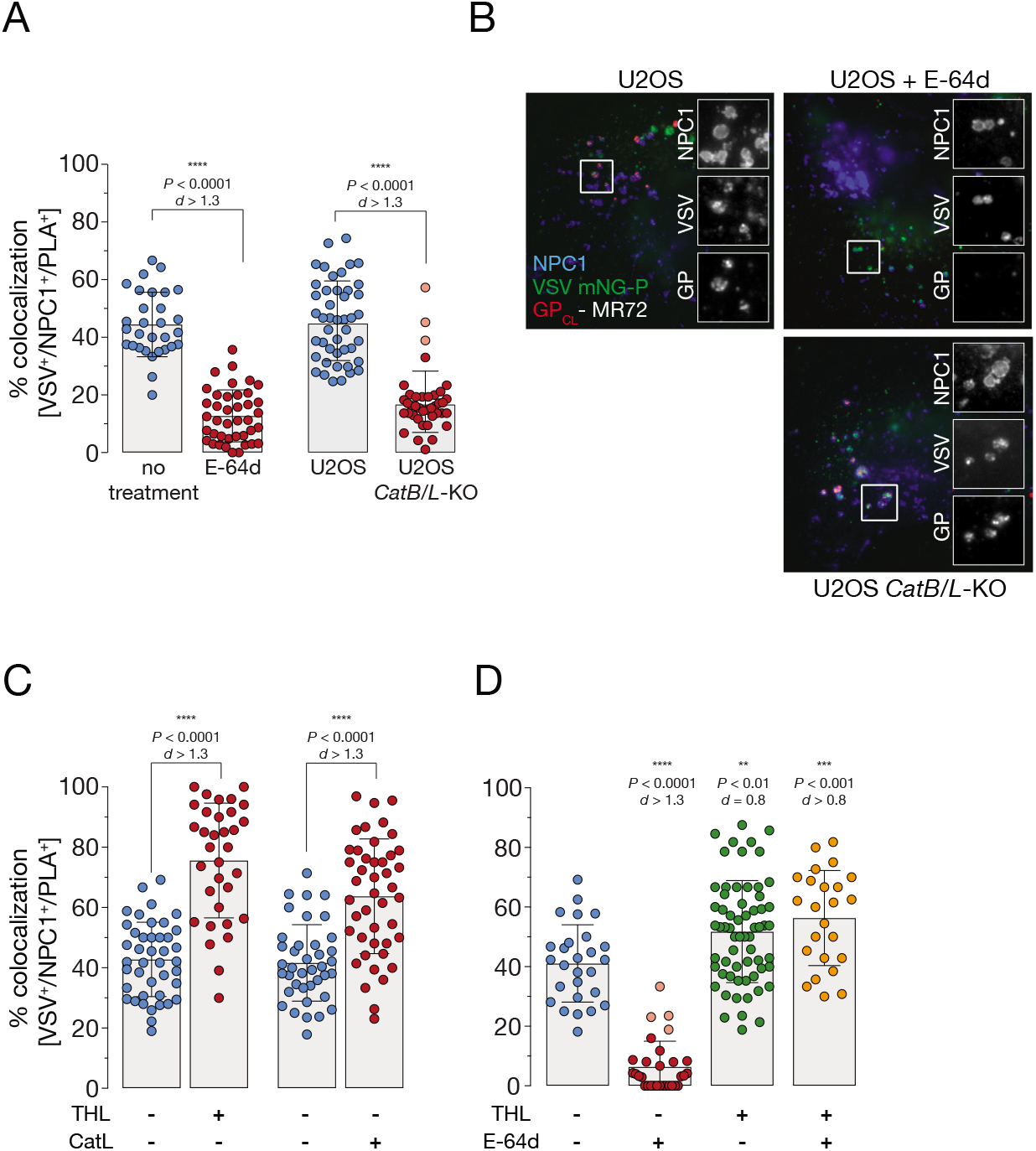
Detection of GP_CL_:NPC1 binding by PLA requires endosomal cleavage of GP. (**A**) VSV mNG-P particles bearing EBOV GP were incubated with either U2OS^NPC1-eBFP2^ cells pre-treated with E-64d (100mM, 6h at 37°C) (left panel), or U2OS *CatB/L-KO* cells ectopically over-expressing flag-tagged NPC1 (right panel). Cells were fixed following virus incubation (1 h at 37°C), permeabilized and subjected to PLA. Data points represent the percentage of VSV^+^/NPC1^+^/PLA^+^ compartments per individual cell; bars depict the average ±SD for all data points pooled from two independent experiments (*n* ≥ 30). Points with reduced transparency represent values outside of the 10th-90th percentile. An unpaired two-tailed t-test was used to compare VSV^+^/NPC1^+^/PLA^+^ compartments of wild-type cells with either inhibitor-treated (left) or KO (right) cells infected by EBOV GP-decorated VSV (****, *P* < 0.0001). Group means calculated from the percentage of VSV^+^/NPC1^+^/PLA^+^ vesicles were compared by Cohen’s *d* effect size. (**B**) Cells described in (A) were exposed to VSV mNG-P EBOV GP uptake for 1h at 37°C. After fixation, viral particles, NPC1 and GP_CL_ were visualized by fluorescence microscopy. Representative images from two independent experiments are shown. (**C**) VSV mNG-P particles bearing EBOV GP were treated *in vitro* with thermolysin (THL) or cathepsin L (CatL), respectively. Viral particles were taken up into U2OS^NPC1-eBFP2^ cells, followed by fixing of the cells and subjecting them to PLA. Data points represent the percentage of VSV^+^/NPC1^+^/PLA^+^ compartments per individual cell; bars depict the average ±SD for all data points pooled from two independent experiments (*n* ≥ 25). To compare VSV^+^/NPC1^+^/PLA^+^ compartments of cells which endocytosed VSV studded with either uncleaved or THL/CatL-cleaved GP, an unpaired two-tailed *t*-test was used (****, *P* < 0.0001). Cohen’s *d* effect size was used to compare the group means calculated from the percentages of VSV^+^/NPC1^+^/PLA^+^ vesicles (*d* > 1.3). (**D**) U2OS^NPC1-eBFP2^ cells were (not) pre-incubated with E-64d (as described in [A]) and VSV mNG-P particles bearing EBOV GP were treated *in vitro* with THL. Following virus internalization into U2OS^NPC1-eBFP2^, cells were fixed and subjected to PLA. Data points represent the percentage of VSV^+^/NPC1^+^/PLA^+^ compartments per individual cell; bars depict the average ±SD for all data points pooled from two independent experiments (*n* ≥ 25). Points with reduced transparency represent values outside of the 10th-90th percentile. To compare VSV^+^/NPC1^+^/PLA^+^ compartments of E-64d-treated cells with those of untreated cells which endocytosed VSV studded with either uncleaved or THL-cleaved GP, an unpaired two-tailed *t*-test was used (**, *P* < 0.01; ***, *P* < 0.001; ****, *P* < 0.0001). Cohen’s *d* effect size was used to compare the group means calculated from the percentages of VSV^+^/NPC1^+^/PLA^+^ vesicles.

GP priming in viral entry can be recapitulated *in vitro* by incubating rVSV bearing full-length EBOV GP with CatL or the bacterial protease thermolysin (THL) which mimics CatL/B cleavage (8); both cleave off GP1 sequences corresponding to the glycan cap and mucin domain. Increased infectivity of viral particles bearing THL/CatL-cleaved GPs was reported previously, and can be attributed to improved cell binding of virions as well as increased accessibility of the GP’s RBS for NPC1 domain C interaction ((7, 11, 25); also see Fig. S3B). *In vitro* cleavage of GP significantly enhanced GP_CL_:NPC1 domain C binding *in situ* by 1.5- to 2-fold indicating that proteolytic processing of GP is indispensable for detection of virus-receptor interaction by proximity ligation (Fig. 3C). In order to determine possible requirements for additional cysteine protease activity pre NPC1-binding, we evaluated proximity ligation with THL-cleaved GP in the presence of the pan-cysteine cathepsin inhibitor E-64d. Here, the absence of cysteine protease proteolytic activity did not affect GP_CL_:NPC1 engagement (Fig. 3D), suggesting the existence of an E-64d-sensitive downstream entry block post-NPC1 binding, as previously proposed (5, 23).

### *In situ* proximity ligation decouples GP_CL_:NPC1 interaction from post-binding entry steps

As one of the critical steps in viral entry, GP:NPC1 interaction is the starting point for subsequent post-binding entry processes including, among others, membrane fusion triggering, cytoplasmic nucleocapsid escape and infection (5, 10–12). To examine if the established *in situ* PLA selectively monitored GP_CL_:NPC1 engagement or, in addition, also downstream steps such as membrane fusion, we made use of an EBOV GP mutant harboring amino acid substitutions in the GP1 internal fusion loop, GP^L529A/I544A^. Amino acids L529 and I544 were shown to form a fusogenic hydrophobic surface at the tip of the fusion loop and proposed to be crucial for insertion of the loop into host membranes and subsequent membrane fusion steps (26, 27). Late-endosomal delivery of VSV bearing GP^L529A/I544A^ and exposure of GP’s RBS did not differ substantially from those of VSV harboring wild-type GP. However, steps post NPC1-binding, including membrane fusion triggering and virus infection were inhibited ((5); also see Fig. S5A). Consistent with these observations, *in situ* proximity ligation of VSV bearing GP^L529A/I544A^ resembled that obtained with VSV decorated with wild-type GP (Fig. 4A).

**Fig. 4.**
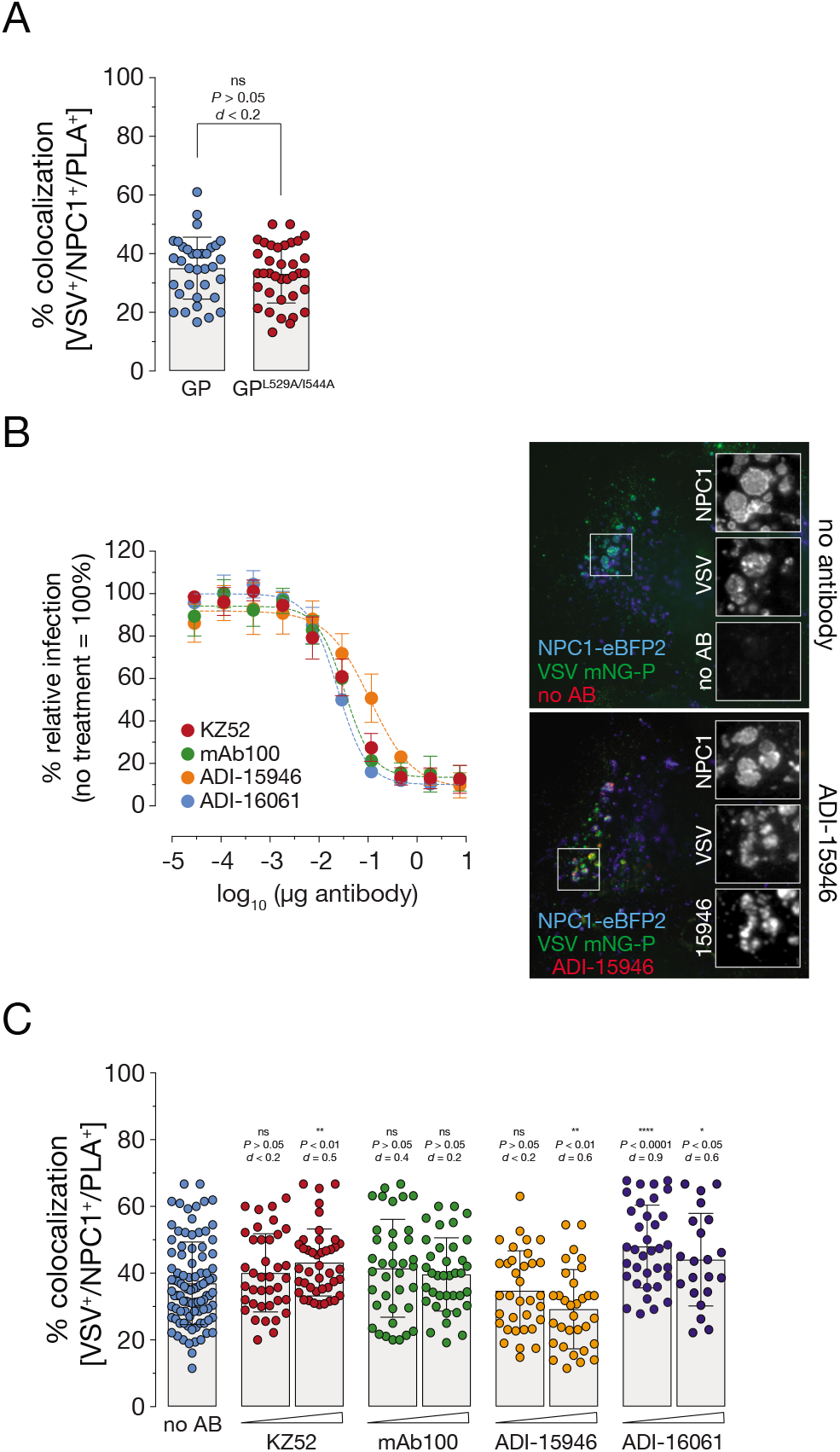
*In situ* proximity ligation decouples GP_CL_:NPC1 interaction from post-binding entry steps. (**A**) VSV mNG-P particles bearing EBOV GP or EBOV GP^L529A/I544A^ were internalized into U2OS^NPC1-eBFP2^ cells for 60 min followed by PLA. Data points represent the percentage of triple positive compartments per individual cell; bars depict the average ±SD for all data points pooled from two independent experiments (*n* ≥ 37). Data analyses included an unpaired two-tailed t-test to compare VSV^+^/NPC1^+^/PLA^+^ vesicles of cells which internalized VSV decorated with wild-type or mutant GP (ns, *P* > 0.05). Cohen’s *d* effect size was used to compare the group means calculated from the percentages of VSV^+^/NPC1^+^/PLA^+^ compartments. (**B**) ADI-15946 was incubated with VSV EBOV GP particles and exposed to U2OS^NPC1-eBFP2^ (right panel). After virus uptake for 1 h at 37°C, cells were fixed and viral particles, NPC1 and bound antibodies were visualized by fluorescence microscopy. Representative images from two independent experiments are shown. Virions were preincubated with increasing amounts of KZ52, mAb100, ADI-15946 or ADI-16061 and then exposed to U2OS^NPC1-eBFP2^ cells for 16 h at 37°C (left panel). Number of infected cells was determined by automated counting of mNG^+^ cells and normalized to infection obtained in the absence of antibodies. Averages ± SD for six technical replicates pooled from two independent experiments are shown. (**C**) VSV mNG-P EBOV GP virions were complexed with KZ52, mAb100, ADI-15946 or ADI-16061 (50 μg/ml and 100 μg/ml) for 1 h at room temperature. Following internalization into U2OS^NPC1-eBFP2^, cells were fixed, and subjected to *in situ* PLA. Data points represent the percentage of VSV^+^/NPC1^+^/PLA^+^ compartments per individual cell; bars show the average ±SD for all data points pooled from two independent experiments (*n* ≥ 22). VSV^+^/NPC1^+^/PLA^+^ vesicles were analyzed by unpaired two-tailed t-test (ns, *P* > 0.05; *, *P* < 0.05; **, *P* < 0.01; ****, *P* < 0.0001) comparing cells which were exposed to virion-antibody complexes to cells exposed to untreated virus. Group means calculated from the percentage of VSV^+^/NPC1^+^/PLA^+^ vesicles were also compared by Cohen’s *d* effect size.

Previous work identified a variety of antibodies with neutralizing activity against EBOV. Those displaying exceptional neutralizing potency were described to target conformational epitopes located in the base subdomain of the GP trimeric complex (28–32). It was proposed that these base-binding antibodies efficiently block fusogenic rearrangements of GP, thereby inhibiting viral entry steps downstream of NPC1 binding. We used these neutralizing antibodies to further investigate if *in situ* PLA monitored GP_CL_:NPC1 binding specifically or also integrated post-receptor binding events (e.g., viral membrane fusion). We tested antibodies targeting the GP1-GP2 interface (KZ52, mAb100), the GP1-GP2 interface plus the glycan cap (ADI-15946), or a subdomain of the fusion loop, the heptad repeat 2 (ADI-16061). Pre-incubation of VSVs with these neutralizing antibodies (50 μg/mL) afforded delivery of virus:antibody complexes to NPC1^+^ LE, but did not hinder virus trafficking to LEs itself or proteolytic processing to GP_CL_ (Fig. 4B, right panel and Fig. S5B-C). However, infection was blocked under these conditions, as expected (Fig. 4B, left panel). Despite their potent neutralizing activity, and consistent with the specified target epitopes as well as previously reported *in vitro* NPC1-binding studies (28–32), base-binding antibodies mAb100 and ADI-15946 had little or no effect on *in situ* PLA, consistent with our hypothesis that this assay selectively detects GP_CL_:NPC1 complex formation (Fig. 4C). Surprisingly, incubation with KZ52 and ADI-16061 slightly improved PLA signal, possibly because they enhanced an upstream step: the delivery of VSV bearing GP_CL_ to NPC1^+^ LE (Fig. 4C).

Collectively, our results provide strong evidence that the *in situ* PLA specifically monitors GP_CL_:NPC1 binding in infected cells and decouples the detection of virus-receptor binding from post-receptor binding steps in viral entry.

### Small molecule inhibitor-mediated selective interference of GP:NPC1 binding delineated by *in situ* PLA

The unprecedented EBOV outbreak in West Africa 2013-16 uncovered a pressing need for anti-EBOV therapeutics; since then, numerous studies screened for and characterized FDA-approved drugs with anti-filoviral activity. The most promising drug candidates, amiodarone, bepridil, clomifene, sertraline, and toremifene were reported to block one or several steps during EBOV entry (16, 33–38). As their precise mechanisms of anti-EBOV activity still remain elusive, we examined the capacity of these inhibitors to alter intracellular GP_CL_:NPC1 interaction in the PLA. As a control, we also tested the well characterized amphiphilic drug U18666A, which was described to block cholesterol export from lysosomes and shown to inhibit EBOV entry at higher concentrations, but not to impact GP_CL_:NPC1 binding *in vitro* (10, 16, 39).

We first titrated the drugs in an EBOV entry assay in wild-type U2OS and U2OS^NPC1-eBFP2^ cells over-expressing NPC1 employed in the PLA (Fig. 5A). Several of the inhibitors were less potent in NPC1-overexpressing cells than in wild-type cells, as reported previously, suggesting they act via the GP_CL_:NPC1 axis ((16); also see Fig. S6A). Some inhibitors hampered VSV trafficking to NPC1^+^ LE, whereas others had little or no effect on viral delivery to NPC1^+^ LE compared to a DMSO-treated control (Fig. S6B). Because these trafficking defects likely contribute to the observed inhibition in infection, we sought to discount these trafficking effects on GP_CL_:NPC1 binding by focusing exclusively on PLA activity in VSV^+^/NPC1 ^+^ compartments. Importantly, these inhibitors exhibited little or no effect on GP cleavage (detected by MR72) or the accessibility of NPC1 domain C (detected by mAb-548; Fig. 5C and Fig. S6C).

**Fig. 5.**
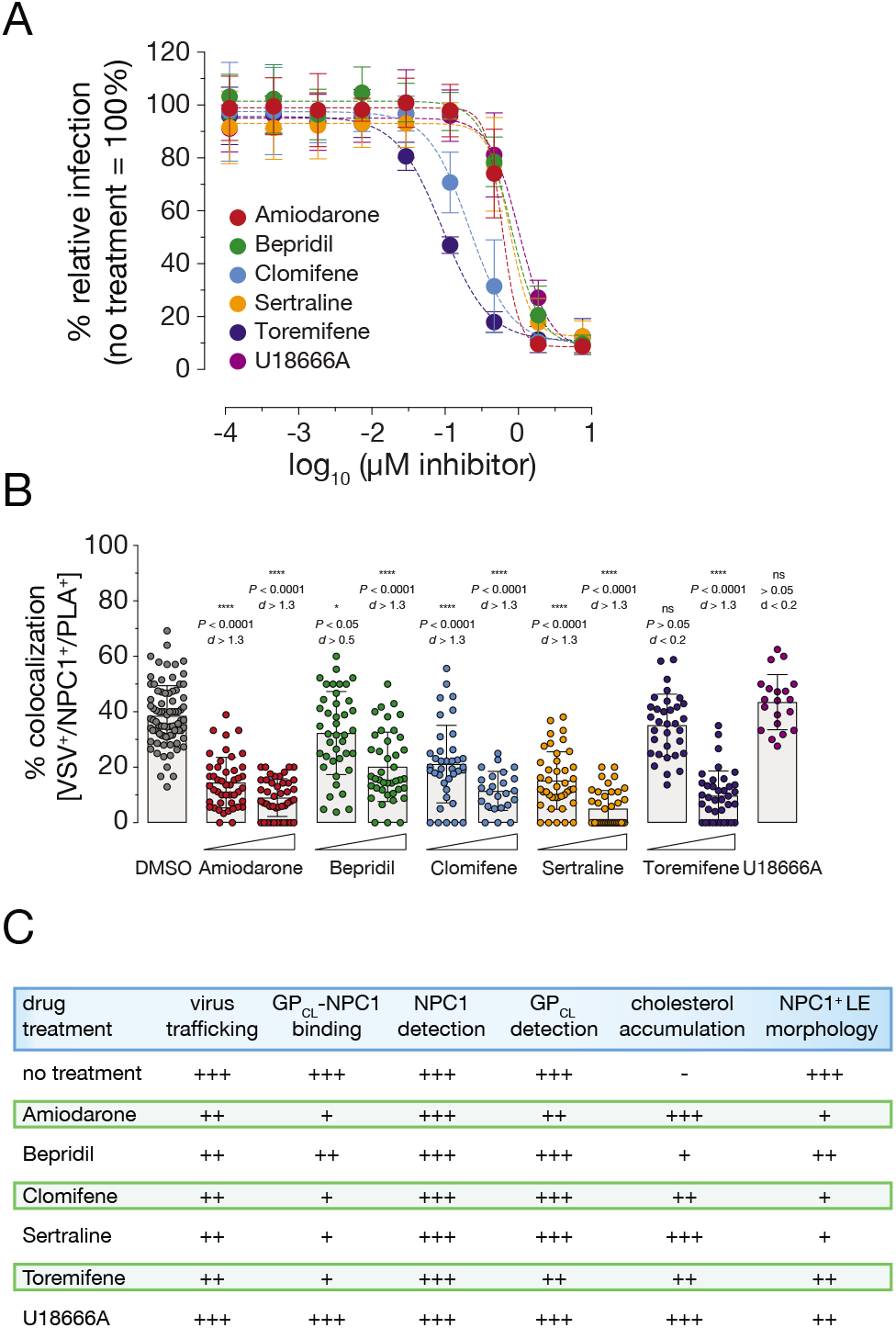
Small molecule inhibitor-mediated selective interference of GP:NPC1 binding delineated by *in situ* PLA. (**A**) U2OS^NPC1-eBFP2^ cells were preincubated with increasing concentrations of amiodarone, bepridil, clomifene, sertraline, toremifene, and U18666A, respectively before exposed to VSV mNG-P EBOV GP for 16 h at 37°C. Infection was measured by automated counting of mNG^+^ cells and normalized to infection obtained in the presence of vehicle only. Averages ± SD for nine to eighteen technical replicates pooled from three to six independent experiments are displayed. (**B**) U2OS^NPC1-eBFP2^ cells were incubated with amiodarone, bepridil, clomifene, and sertraline, respectively (2 μM or 5 μM, 1 h at 37°C), or toremifene (2 μM or 10 μM, 1 h at 37°C), or U18666A (10 μM, 2 h at 37°C), followed by VSV mNG-P EBOV GP uptake for 1 h. Cells were subjected to *in situ* PLA and analyzed by fluorescence microscopy. The percentage of triple positive VSV^+^/NPC1^+^/PLA^+^ compartments per individual cell is represented by data points; bars show the average ±SD for all data points pooled from two independent experiments (*n* ≥ 20). VSV^+^/NPC1^+^/PLA^+^ vesicles were analyzed by unpaired two-tailed *t*-test (ns, *P* > 0.05; *, *P* < 0.05; ****, *P* < 0.0001) comparing inhibitor-exposed to untreated cells. Group means calculated from the percentage of VSV^+^/NPC1^+^/PLA^+^ compartments were also compared by Cohen’s *d* effect size. (**C**) Table summarizing results from Fig. 5B and Fig. S6.

Screening the inhibitory agents via PLA revealed that amiodarone, bepridil, clomifene, sertraline, and toremifene all significantly interfered with GP_CL_:NPC1 interaction as compared to a DMSO-treated control (Fig. 5B). By contrast, but consistent with previous *in vitro* data, U18666A did not block GP_CL_:NPC1 binding (Fig. 5B). The drug treatments also substantially modified NPC1 trafficking to LE and/or LE localization: we observed pronounced imbalances in NPC1 distribution, illustrated by LE vesicles with extensively elevated or decreased NPC1 levels accumulating in the perinuclear regions of cells (Fig. 5C and S6C). Further, NPC1’s transporter activity was strongly impaired: although cells did not exhibit cholesterol accumulation in NPC1^+^ LE following the short-term drug treatments carried out prior to PLA measurements, prolonged inhibitor incubations (16 h instead of 1 h incubation prior to PLA) clearly blocked cholesterol clearance from LE (Fig. 5C and Fig. S6C, upper panel).

In summary, our results imply that a number of FDA-approved small molecule inhibitors interfere with GP_CL_:NPC1 engagement, and such block downstream viral entry steps. However, our findings and previously published work indicate that the inhibitory mode of action of these compounds is likely multifactorial. Although our data suggests that some of the compounds directly affect GP_CL_:NPC1 interaction in endosomal compartments during entry, they also uncover contributions from drug-induced changes in the morphology and distribution of NPC1^+^ compartments.

## DISCUSSION

Interaction of the filovirus spike protein GP with its universal intracellular receptor NPC1 is indispensable for viral entry and infection (10–14). However, our understanding of the mechanism of GP:NPC1 recognition is largely derived from *in vitro* assays that use detergent-extracted NPC1 or a soluble form of the GP-interacting domain C of NPC1 and *in vitro*-cleaved GP_CL_ (11, 12). Although powerful, these assays likely do not fully recapitulate the authentic virus-receptor interaction in late endo/lysosomal compartments. As a case in point, a number of small molecule entry inhibitors that appear to target NPC1 in cells do not block GP_CL_:NPC1 domain C interaction *in vitro*, leaving questions about their mechanism of action open (16).

Here, we describe a novel assay that monitors the GP:NPC1 interaction in intact cells. Detection of the virus-receptor complexes *in situ* is based on the principle of DNA-assisted, antibody-mediated proximity ligation (20, 21). Several studies have used PLAs to detect interaction of viruses with host proteins at the plasma membrane (40), in the cytoplasm (41, 42) and early endosomes (43). To our knowledge, our assay represents the first example to afford visualization of protein-protein interactions in late endosomal/lysosomal compartments.

Previous studies used *in vitro* protease treatments in conjunction with infectivity measurements in protease inhibitor-treated or genetically engineered cells to establish a requirement for GP cleavage in EBOV entry (7–9). Further, *in vitro* binding studies with soluble NPC1 domains demonstrated that GP cleavage is required for virus-receptor interaction (10–14). Here, we used the PLA to directly investigate the role of GP proteolytic cleavage in GP:NPC1 interaction within cellular endo/lysosomal compartments. Concordant with our current understanding, we found that blockade of all endosomal cysteine cathepsins with the pan-cysteine cathepsin inhibitor E-64d abrogated both GP cleavage and GP:NPC1 association *in situ* and that *in vitro* pre-cleavage of EBOV GP to GP_CL_ significantly boosted this interaction. We also observed that genetic knockout of the two key cysteine cathepsins identified previously–CatB and CatL–substantially inhibited GP:NPC1 association as measured by PLA. Unexpectedly, knocking out *CatB* and *CatL* did not prevent exposure of the RBS in endosomes, as judged by the continued capacity of RBS-specific mAb MR72 to detect viral particles. Thus, one or more non-CatB/CatL cysteine cathepsins appear to at least partially cleave GP in endosomes in a manner that, however, does not permit stable GP:NPC1 association. Alternatively, it is possible, at least in principle, that CatB and/or CatL mediate additional yet undiscovered cleavage events that are required for GP:NPC1 binding.

Similar to published *in vitro* GP:NPC1 binding assays, our *in situ* PLA also specifically monitored GP:NPC1 interactions decoupling them from downstream post-binding steps allowing us to entirely focus on molecular mechanisms underlying the indispensable receptor binding. We employed fusion-blocking GP mutants and neutralizing antibodies targeting epitopes essential for GP’s conformational rearrangements during membrane fusion; both interventions however left GP_CL_:NPC1 binding measured by PLA largely unaffected. Previous publications suggested a dual mechanism of action for several of these antibodies, KZ52, mAb100, and ADI-15946 (28, 30); to block EBOV GP-mediated infection they hampered post-NPC1-binding steps, and interfered with GP’s proteolytic processing *in vitro*. Surprisingly, the latter *in vitro* findings were not recapitulated by the *in situ* PLA. Our results suggest that in the presence of neutralizing antibodies the Cat B/L cleavage sites located in GP’s β13-14 loop were readily accessible in an *in situ* endosomal environment while they did not appear to be in the *in vitro* cell-free system. While the RBS-specific mAb MR72 does not allow to monitor minor changes in GP cleavage efficiency, our PLA results indicate sufficient cleavage to allow efficient NPC1 binding. Further, our experiments suggest that *in vitro* cleavage studies do not adequately recapitulate the authentic interactions within the trifecta of endosomal proteases, virus glycoproteins and GP-targeting mAbs in an acidic endosomal environment.

Lastly, by examining the effect of small molecule inhibitors on GP_CL_:NPC1 binding, we further unraveled their multifaceted mechanisms of inhibitory action towards EBOV GP-mediated infection. All FDA-approved drugs tested, amiodarone, bepridil, clomifene, sertraline, and toremifene, significantly hampered GP_CL_:NPC1 interaction measured by *in situ* PLA. Previous reports focused on bepridil, sertraline and toremifene destabilizing the GP’s prefusion conformation by binding to a cavity at the interface of GP1 and GP2 *in silico* (35, 38). Our data also indicated striking changes in the intracellular distribution and morphology of NPC1^+^ LE/LY, as well as NPC1’s cholesterol transporter function which was most notable for toremifene, clomifene, and sertraline suggesting they (in addition) also acted via the GP_CL_:NPC1 axis; overexpression of NPC1 clearly counteracted their inhibitory mechanism during infection. We propose that the loss of GP_CL_:NPC1 binding is most likely caused by adverse effects on both, NPC1 and GP, leading to destabilization of GP and drastic modifications in the NPC1^+^ LE/LY phenotype. Resolving the effects of small molecule inhibitors on GP_CL_:NPC1 binding remains the focus of ongoing work and will help us to decode this interaction in more detail.

In summary, the newly established *in situ* proximity ligation assay represents a powerful tool to delineate molecular mechanisms underlying receptor-filoviral glycoprotein interactions, to characterize host factors modulating EBOV entry, and to unravel the mode of action of antibodies and small molecule inhibitors in a cell-based system. Our data indicates that *in vitro* and *in silico* studies, while informative, have severe limitations in adequately recapitulating authentic receptor-glycoprotein interactions in host cells. Hence, we propose to modify and translate the employed *in situ* assay to unravel the receptor interactions of diverse viral surface proteins.

## MATERIAL AND METHODS

### Cells and viruses

Human osteosarcoma U2OS cells were cultured in modified McCoy’s 5A medium (Life Technologies) supplemented with 10% fetal bovine serum (Atlanta Biologicals), 1% penicillin-streptomycin (Life Technologies), and 1% GlutaMax (Life Technologies). Cells were kept at 37°C with 5% CO2 in a humidified incubator. U2OS cells stably overexpressing NPC1-eBFP2 were generated as described previously and maintained under the same conditions mentioned above (17). Propagation of recombinant vesicular stomatitis Indiana virus (rVSV) expressing eGFP in the first position and bearing the VSV G or EBOV GP glycoprotein, derived from EBOV/H.sap/COD/76/Yam-Mayinga (EBOV “Mayinga” isolate) as well as an rVSV bearing the EBOV GP glycoprotein, and an mNeongreen-phosphoprotein P (mNG-P) fusion protein has been described previously (5, 44–46). Pseudotyped VSVs bearing mNG-P and variant GPs, GP^T83M/K114E/K115E^ and GP^L529A/I544A^ (both also lacking the mucin-like domain), were prepared as previously reported (7, 24, 47).

For some experiments, cleaved viral particles bearing GP_CL_ were first generated by incubation with thermolysin (THL, 1 mg/ml, pH 7.5, 37°C for 1h; Sigma) or recombinant human cathepsin L (CatL, 2 ng/ul, pH 5.5, 37°C for 1h; R&D Systems) as described previously (5). Reactions were stopped by removal onto ice and addition of phosphoramidon (1 mM; Peptides International) or E-64 (10 μm; Peptides International), respectively. While viral particles cleaved with CatL were used immediately, THL-cleaved virus was kept at −80°C until usage.

### Antibodies

For immunofluorescence analysis, NPC1-eBFP2 was detected by a rabbit anti-BFP antibody (GeneTex) followed by a secondary anti-rabbit antibody-Alexa 405 or -Alexa 488 fluorophore (Thermo Scientific). During proximity ligation assay (PLA), NPC1 was detected by mAb-548, whose generation was described earlier (17). Detection of proteolytically cleaved GP was carried out with a RBS-specific human anti GP antibody, MR72, either followed by a secondary anti-human antibody-Alexa 555 fluorophore or following the PLA protocol as described below. To detect EBOV GP by immunofluorescence analysis and to determine the effect of GP-targeting antibodies on GP_CL_:NPC1 interaction we used the previously described human anti-GP antibodies KZ52, mAb100, ADI-15946, and ADI-16061 (28–32). For production of KZ52, ADI-15946, and ADI-16061, variable heavy- and light-chain domain sequences were cloned into the mammalian expression vectors pMAZ-IgL (encoding for the expression cassette of human κ light-chain constant domains) and pMAZ-IgH (encoding for the expression cassette of human γ1 chain constant domains). Antibody production in FreeStyle™ 293-F cells (ThermoFisher) and subsequent purification were carried out as described earlier (17).

### Inhibitors

Stock solutions of drugs were prepared in dimethyl sulfoxide (DMSO) and stored as frozen aliquots until use. Cells were incubated with 3.47 (Microbiotix), Amiodarone (Sigma), Bepridil (Sigma), Clomifene (Sigma), Sertraline (Toronto Research Chemicals), or Toremifene (Sigma) for 1 h or 16 h at 37°C with concentrations as indicated. Incubation of cells with E-64d (Peptides International) was extended to 6 h at 37°C, while cells were not preincubated with U18666A (Calbiochem) for PLA, but preincubated for 1 h or 16 h at 37°C for additional described experiments at concentrations indicated. Inhibitors were maintained at the same concentrations during virus spin-oculation and virus entry into cells.

### Generation of U2OS *CatB/L* double knockout cells overexpressing NPC1

U2OS cells were transduced with a lentivirus carrying human codon-optimized *Streptomyces pyogenes* Cas9 (spCas9) and blasticidin resistance genes to generate U2OS-Cas9 cells expressing Cas9. Briefly, 293FT cells were co-transfected with Cas9-expressing plasmid lentiCas9-Blast (Addgene #52962, a gift from Feng Zhang), the lentiviral packaging plasmid psPAX2 (Addgene #12260, a gift from Didier Trono) and a VSV G expressing plasmid. The supernatant filtered through a 0.45 μm filter was used to transduce U2OS cells in the presence of 6 μg/ml of polybrene and transduced cells were selected with 15 μg/ml of blasticidin. A lentiGuide-Puro plasmid (Addgene plasmid #52963, a gift from Feng Zhang) expressing human *CatL*-targeting sgRNA (5’-CTTAGGGATGTCCACAAAGC-3’, targets anti-sense strand, nt 1021-1040 of the *CatL* transcript variant 1 mRNA or nt 4205-4186 of *CatL* gene ID 1514) was used to generate lentiviruses as described above. U2OS-Cas9 cells transduced with these lentiviruses were selected with 2 μg/ml of puromycin. Editing of the *CatL* gene was confirmed by Sanger sequencing of a 673-bp amplicon using primers flanking the sgRNA target site. A single cell clone carrying a homozygous deletion of 15 nucleotides that is predicted to disrupt a critical N-glycosylation site (204-NDT-206) that is required for lysosomal targeting of CatL by causing a T206I change (along with a deletion of aa 207-210) was selected (48, 49). Absence of wild-type allele from the *CatL* knockout (KO) cells was confirmed by RT-PCR using primers specific for the 15-nt deletion. To generate the *CatB/L* double KO, the *CatL*-KO cells were further transduced with a lentivirus encoding *CatB*-specific sgRNA (5’-TTGACCAGCTCATCCGACAG-3’, targets anti-sense strand, nt 251-270 of human *CatB* transcript variant 1 mRNA or nt 14,791-14,772 of human *CatB* gene ID 1508). A single cell clone carrying an insertion of a single nucleotide (T) that is predicted to cause a frame-shift leading to a truncated pro-peptide of 28 amino acids (frame shift at aa position 27) was selected. Absence of *CatB* and *CatL* activity in this single cell clone was confirmed by cathepsin activity assays (described below). Stable U2OS cells expressing NPC1 C-terminally tagged with a triple flag sequence in the *CatB/L*-KO background were generated as described earlier. In short, retroviruses packaging the transgene were produced by triple transfection of 293T cells, and target U2OS cells were directly exposed to sterile-filtered retrovirus-laden supernatants in the presence of polybrene (6 μg/ml). Transduced cell populations were selected with puromycin (2 μg/ml) and expression of flag-tagged NPC1 was confirmed by immunostaining with an anti-Flag M2 antibody (Sigma).

### Cathepsin L and B activity assay

U2OS-Cas9, U2OS *CatL*-KO, U2OS *CatB*-KO, and U2OS *CatB/L-KO* cells stably expressing flag-tagged NPC1 were lysed (50 mM MES [pH 5.5], 135 mM NaCl, 2 mM ethylenediaminetetraacetic acid [EDTA], 0.5% Triton X-100) for 1 h on ice. As a control, cleared U2OS-Cas9 lysates were pre-incubated with the protease inhibitor E-64 (20 μM; Peptides International) for 20 min at room temperature (when indicated). Then, lysates were mixed with reaction buffer (100 mM NaAcetate [pH 5], 1 mM EDTA, 4 mM dithiothreitol), incubated with the fluorogenic peptide substrate Z-FR-AMC (150 μM; R&D Systems) and measured at a fluorometer following a 0 to 30 min incubation (ƛ_Ex_ = 390 nm, ƛ_Em_ = 460 nm). Measurements of substrate hydrolysis were normalized to maximum hydrolysis on U2OS-Cas9 cells reached after 30 min; data represents the mean value and standard deviations of three independent experiments (*n* = 3).

### Infection experiments

Confluent U2OS cells were infected with pre-titrated amounts of pseudotyped VSV particles bearing wild-type or mutant GPs. Prior to infection, VSVs were diluted in corresponding media, and infected cells were maintained at 37°C for 14 to 16 h post infection before manual counting of eGFP^+^ and mNG^+^ cells or automated counting using a Cytation 5 cell imaging multi-mode reader (BioTek Instruments) and a CellInsight CX5 imager (Thermo Fisher) including onboard software. When indicated, cells were pre-incubated with small molecule inhibitors diluted in corresponding media; for antibody neutralization experiments, VSV particles were incubated with increasing concentrations of test Ab at room temperature for 1 h, prior to addition to cell monolayers. Virus infectivities were measured as described above. Virus neutralization data was subjected to nonlinear regression analysis (4-parameter, variable slope sigmoidal dose-response equation; GraphPad Prism).

### Immunofluorescence microscopy and immunostaining

To investigate determinants of VSV mNG-P EBOV GP internalization into endosomal compartments, pre-titrated amounts of VSV particles bearing mNG-P and either wild-type or mutant EBOV GP were diluted into imaging buffer (20 mM HEPES [pH 7.4], 140 mM NaCl, 2.5 mM KCl, 1.8 mM CaCl_2_, 1 mM MgCl_2_, 5 mM glucose, 2% FBS), and spin-occulated onto pre-chilled U2OS^NPC1-eBFP2^ cells on coverslips. Unbound virus was removed by washing with cold PBS. Cells were then placed in warm imaging buffer, and allowed to internalize VSVs for 1 h at 37°C. Cells were fixed with 3.7% paraformaldehyde and permeabilized with PBS/0.1% Triton X-100. NPC1-eBFP2 was detected with either primary mouse anti-NPC1 mAb-548 or rabbit anti-BFP antibodies and secondary anti-mouse or anti-rabbit antibody-Alexa 405 fluorophore conjugates, respectively (Thermo Scientific). Cleaved GP was detected with the primary mouse RBS-specific anti-GP antibody MR72 followed by secondary anti-mouse antibody-Alexa 555 fluorophore conjugate (Thermo Scientific). To investigate transport of IgGs bound to VSV particles into NPC1^+^ endosomal compartments, VSV particles bearing mNG-P were preincubated with antibodies (50 or 100 μg/ml, respectively) for 1 h at room temperature, followed by internalization into target cells and detection by incubation with secondary anti-human-Alexa 555 fluorophore conjugates (Thermo Scientific). Cells were examined by immunofluorescence analysis performed on an Axio Observer Z1 widefield epifluorescence microscope (Zeiss Inc.) equipped with an ORCA-Flash4.0 LT digital CMOS camera (Hamamatsu Photonics), a 63x/1.4 numerical aperture oil immersion objective and a DAPI (4’,6-diamidino-2-phenylindole)/fluorescein isothiocyanate (FITC)/tetramethyl rhodamine isocyanate (TRITC)/Cy5 filter set. Images were processed in Photoshop (Adobe Systems).

### Cholesterol accumulation assay

Cholesterol accumulation following inhibitor treatment of U2OS^NPC1-eBFP2^ cells was monitored by incubation with filipin (50 μg/ml; Sigma) for 1 h at room temperature. Cells were examined by immunofluorescence analysis as described under the method section ‘Immunofluorescence microscopy and immunostaining’.

### Proximity Ligation Assay

U2OS^NPC1-eBFP2^ cells were allowed to internalize VSV particles (in the presence or absence of small molecule inhibitors or antibodies), fixed and permeabilized as described in the method section ‘Immunofluorescence microscopy and immunostaining’. MR72, a RBS-targeting anti-GP mAb and mAb-548, an NPC1 domain C-targeting mAb were directly labeled with Duolink^®^ In Situ Probemaker PLUS and Duolink^®^ In Situ Probemaker MINUS (Sigma) following the manufacturer’s instructions. Fixed cells were incubated with labeled antibodies in a humidity chamber at 37°C for 1 h. Excess antibody was removed by washing with Duolink^®^ In Situ Wash Buffer (Sigma). GP:NPC1 interaction was detected by applying the Duolink^®^ In Situ Detection Reagents Red kit following the manufacturer’s instructions (Sigma). After removing excess reagents, NPC1-eBFP2 was detected using a rabbit anti-BFP antibody followed by a secondary anti-rabbit antibody-Alexa 405 fluorophore conjugate (Thermo Fisher). Cells were examined by immunofluorescence analysis as described under the method section ‘Immunofluorescence microscopy and immunostaining’.

## ACKNOWLEDGMENTS

We acknowledge Isabel Gutierrez, Estefania Valencia, Laura Polanco, and Cecelia Harold for laboratory management and technical support. This work was supported by National Institutes of Health (NIH) grant R01AI134824 (to K.C.). K.C. is a member of the Scientific Advisory Board of Integrum Scientific, LLC.

---

**Fig. S1.** PLA requires the presence of both detecting antibodies and VSV internalization. VSV mNG-P particles bearing EBOV GP were exposed to U2OS^NPC1-eBFP2^ cells for 1 h at 37°C or, to inhibit endocytosis, at 4°C. Cells were fixed, permeabilized and subjected to *in situ* PLA using GP_CL_-specific (MR72) or NPC1 domain C-specific (mAb-548) antibodies only, or a combination of both. The percentage of VSV^+^/NPC1^+^/PLA^+^ compartments per cell was determined by fluorescence microscopy and presented here by individual data points. Graphic bars show the average ±SD for all data points pooled from one to two independent experiments (*n* ≥ 10).

**Fig. S2.** GP_CL_:NPC1 interface formation is required for *in situ* PLA. (**A**) VSV particles bearing EBOV GP or EBOV GP^T83M/K114E/K115E^ were exposed to U2OS^NPC1-eBFP2^ cells for 1 h at 37°C. After fixation, viral particles, NPC1 and GP_CL_ were visualized by fluorescence microscopy. Representative images from two independent experiments are shown. (**B**) Virions bearing EBOV GP or EBOV GP^T83M/K114E/K115E^ normalized for the VSV matrix protein M (data not shown) were used to infect U2OS^NPC1-eBFP2^ cells. Infection was measured by manual counting of mNG^+^ cells and normalized to infection with VSV mNG-P decorated with wild-type GP. Averages ± SD for four technical replicates pooled from two independent experiments are presented. (**C**) After incubation of U2OS^NPC1-eBFP2^ cells with 3.47 (1 μM, 1 h at 37°C), cells were either directly fixed or exposed to VSV mNG-P EBOV GP for 1 h at 37°C prior to fixation. NPC1 was detected by mAb-548 (upper panel) and GP_CL_ was detected by MR72 (lower panel). Representative images of fluorescence microscopy from two independent experiments are shown.

**Fig. S3.** Endosomal protease activity is essential for EBOV GP-mediated particle infectivity. (**A**) U2OS^NPC1-eBFP2^ cells were treated with E-64d (100 mM, 6 h at 37°C) followed by infection with VSV bearing EBOV GP. Number of infected cells was determined by manual counting of mNG^+^ cells and normalized to infection obtained in the presence of vehicle only. Averages ± SD for four technical replicates pooled from two independent experiments are presented. (**B**) After *in vitro* treatment of VSV mNG-P EBOV GP particles with thermolysin (THL) or cathepsin L (CatL), virions were normalized by a quantitative Western Blot assay detecting VSV M and exposed to U2OS^NPC1-eBFP2^ cells. Number of infected cells was determined by manual counting of mNG^+^ cells and normalized to infection obtained with untreated VSV mNG-P EBOV GP. Averages ± SD for four technical replicates pooled from two independent experiments are presented.

**Fig. S4.** Characterization of CRISPR/Cas9-generated U2OS *CatB/L-KO* cells. (**A**) Alignment of the wild-type *Cat L* and *B* gene sequences with alleles in the *CatB/L*-KO cell clone. The gRNA target sequence is depicted in red, the PAM sequence is depicted in blue. (**B**) *CatB* and *CatL* activities in U2OS cell extracts were measured by fluorogenic peptide turnover. As a control, the proteolytic activity in U2OS *CatL*- or *CatB*-KO cells and U2OS cells which were pretreated for 20min with 20 μM E-64d was also determined. Averages ± SD for six technical replicates pooled from two independent experiments are presented. (**C**) Susceptibility of U2OS *CatB/L*-KO cells and U2OS *CatB/L*-KO cells ectopically overexpressing flag-tagged NPC1 to EBOV GP and VSV G-mediated VSV infection. Number of infected cells was determined by manual counting of eGFP^+^ or mNG^+^ cells and normalized to infection obtained with wild-type U2OS cells.

**Fig. S5.** *In situ* PLA allows decoupling of GP:NPC1 binding from post-NPC1 binding steps. (**A**) VSV mNG-P particles studded with EBOV GP or EBOV GP^L529A/I544A^ were internalized into U2OS^NPC1-eBFP2^ cells for 1 h. Following fixation, viral particles, NPC1 and GP_CL_ were visualized by fluorescence microscopy. Representative images from two independent experiments are shown. (**B**) Virions bearing EBOV GP were first complexed with KZ52, mAb100 or ADI-16061 antibodies (50μg/ml) for 1 h at room temperature, and then virus-antibody complexes were internalized into U2OS^NPC1-eBFP2^ cells. By fluorescence microscopy, viral particles, NPC1 and bound antibodies were visualized. (**C**) Samples were generated as described in (B) and viral particles, NPC1 and GP_CL_ (via MR72) were visualized by fluorescence microscopy.

**Fig. S6.** Small molecule inhibitors interfere with cell susceptibility to infection, virus trafficking as well as NPC1 distribution and function. (**A**) Wild-type U2OS or U2OS^NPC1-eBFP2^ cells were preincubated with increasing amounts of indicated inhibitors and then exposed to VSV decorated with EBOV GP for 16 h at 37°C in presence of the inhibitors. Number of infected cells was determined by automated counting of eGFP^+^ cells and normalized to infection obtained in the absence of inhibitors. Averages for six technical replicates pooled from two independent experiments are shown. (**B**) As outlined in Fig. 5B, U2OS^NPC1-eBFP2^ cells were incubated with amiodarone, bepridil, clomifene, toremifene, sertraline, and U18666A respectively, followed by VSV mNG-P EBOV GP uptake for 1 h. Cells were subjected to *in situ* PLA and analyzed by fluorescence microscopy. The percentage of double positive VSV^+^/NPC1^+^ compartments per individual cell is represented by data points; bars show the average ±SD for all data points pooled from two independent experiments (n ≥ 20). VSV^+^/NPC1 ^+^ vesicles were analyzed by unpaired two-tailed *t*-test (ns, *P* > 0.05; *, *P* < 0.05; ***, *P* < 0.001; ****, *P* < 0.0001) comparing inhibitor-exposed to vehicle treated cells. (**C**) U2OS^NPC1-eBFP2^ cells were incubated with U18666A (10μM), amiodarone (5μM), clomifene (5μM) or DMSO carrier for 16 h at 37°C, followed by detection of NPC1 by mAb-548 (bottom panels) or NPC1 detection followed by filipin staining highlighting cholesterol accumulations (top panels) and analyzed by fluorescence microscopy.

